# The phylodynamic threshold of measurably evolving populations

**DOI:** 10.1101/2025.07.07.663495

**Authors:** Ariane Weber, Julia Kende, Camila Duitama González, Sanni Översti, Sebastian Duchene

## Abstract

The molecular clock is a fundamental tool for understanding the time and pace of evolution, requiring calibration information alongside molecular data. Sampling times are often used for calibration since some organisms accumulate enough mutations over the course of their sampling period. This practice ties together two key concepts: measurably evolving populations and the phylodynamic threshold. Our current understanding suggests that populations meeting these criteria are suitable for molecular clock calibration via sampling times. However, the definitions and implications of these concepts remain unclear. Using Hepatitis B virus-like simulations and analyses of empirical data, this study shows that determining whether a population is measurably evolving or has reached the phylodynamic threshold does not only depend on the data, but also on model assumptions and sampling strategies. In Bayesian applications, a lack of temporal signal due to a narrow sampling window results in a prior that is overly informative relative to the data, such that a prior that is potentially misleading typically requires a wider sampling window than one that is reasonable. In our analyses we demonstrate that assessing prior sensitivity is more important than the outcome of tests of temporal signal. Our results offer guidelines to improve molecular clock inferences and highlight limitations in molecular sequence sampling procedures.

## 1 Introduction

Molecular sequence data have become nearly ubiquitous for studying the evolution of modern and ancient organisms. A fundamental concept in molecular evolution is the ‘molecular clock’, which posits that substitutions accumulate roughly constantly over time (Zuckerkandl and Pauling, 1965). An underlying assumption of the classic molecular clock is that selective constraints are negligible for most sites and over time. The development of molecular clock models as statistical processes relaxes this and other assumptions by allowing for rate variation among branches (and sometimes sites, see Ho 2014) in phylogenetic trees (reviewed by Guindon (2020), Ho and Duchêne (2014)).

Molecular clock models necessarily involve two key quantities, the evolutionary timescale and the ‘evolu tionary rate’, with the latter representing the combination of mutations and substitutions that accrue over time. However, evolutionary times and rates are unidentifiable (Dos Reis and Yang (2013), as reviewed by Bromham et al. 2018, Guindon 2020), and therefore cannot be jointly estimated using genetic sequence data alone. To make inferences from genetic sequences, all molecular clock methods require prior assumptions about evolutionary times or rates, known as a ‘molecular clock calibration’. Three main calibrations exist: First, the age of the most recent common ancestor between two samples can be constrained to a given time point or interval, known as an ‘internal node calibration’. Second, a known estimate of the evolutionary rate can be incorporated (e.g. as a prior in Bayesian frameworks). Third, in cases where molecular sequences are sampled at different points in time (heterochronous sampling), the tips of the phylogeny can be anchored to these time points (‘ tip-calibration’; reviewed by Rieux and Balloux (2016)). The choice of calibration depends on the information available and its reliability (Duchêne et al., 2014, Warnock et al., 2012). For instance, it would be remiss to ignore evidence about when two lineages shared a common ancestor if the fossil record is compelling (Gavryushkina et al., 2017, Ronquist et al., 2016). Crucially, multiple sources of calibration information can be provided for the molecular clock.

### 1.1 Measurably evolving populations

Rapidly evolving organisms, notably many viruses and bacteria, have been found to accrue an appreciable number of substitutions over the sampling timescale. Influenza viruses, for example, have evolutionary rates of around 6 × 10^−3^ subs/site/year (substitutions per genomic site per year) (Ghafari et al., 2022, Sanjuán, 2012). Assuming a genome size of 13,500 nucleotides, one would expect to observe one mutation every 4 to 5 days 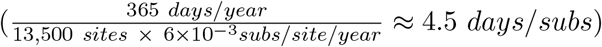. If genome samples are collected over the course of a few weeks, the sampling times themselves can be used to calibrate the molecular clock, and tip-calibration is therefore warranted. Data sets for which tip-calibration is feasible are considered to have been sampled from a ‘measurably evolving population’ (Drummond et al., 2003b) and to have ‘temporal signal’.

Measurably evolving populations are typically characterised either by a sampling period that is long relative to the evolutionary rate, a sufficiently big data set (long molecular sequences or many samples), or both. Traditionally, such characteristics were mainly found in rapidly evolving organisms, typically RNA viruses. Nowadays, advances in sequencing technologies have dramatically expanded the range of organisms from which data sets can be considered to have been sampled from a measurably evolving population. Namely, ancient DNA techniques have effectively expanded the genome sampling window for many organisms (Duchene et al., 2020b, Spyrou et al., 2019a), and whole genome sequencing has meant that data sets of ‘slowly’ evolving microbes often carry sufficient information for calibrating the molecular clock (Biek et al., 2015) even when the sampling period covers only a few decades (Menardo et al., 2019).

### 1.2 The phylodynamic threshold

Genomic data sets collected during the early stages of an outbreak, for example, often pose two problems: low genetic diversity and a narrow sampling window. Both can lead to highly uncertain estimates of evolutionary rate and time of origin. The point in time when an organism has accumulated sufficient genetic changes since its emergence to allow for informative tip-calibration is referred to as the ‘phylodynamic threshold’ (Duchene et al., 2020a). At a minimum, tip-calibration requires that one mutation has occurred over the sampling period for the method to be informative. For a given organism, the minimum sampling period can be calculated as the inverse of the product of genome size and the evolutionary rate (i.e. 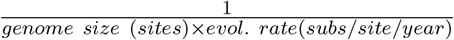 = years to observe one mutation). We refer to this amount of time as the expected phylodynamic threshold.

The terms phylodynamic threshold and measurably evolving population are different, albeit related, concepts. A population is measurably evolving if the samples available are sufficiently informative as to allow for tip-calibration. In contrast, the phylodynamic threshold is the amount of time over which we would need to draw samples after their emergence for them to behave as if being sampled from a measurably evolving population. For a recently evolving pathogen the phylodynamic threshold would simply correspond to the time until it can be considered a measurably evolving population, under the condition that data have been collected since the emergence of the organism. In contrast, an organism that emerged further in the past may have accumulated considerable genetic diversity over time, effectively reaching its phylodynamic threshold, but if samples are drawn from a very short time window they may fail to capture a representative amount of such genetic diversity.

### 1.3 Tests of temporal signal

Our ability to extract information from a tip-calibration framework can be assessed through tests of temporal signal. The importance of performing such tests arises from the observation that a lack of temporal signal is associated with unreliable evolutionary rate estimates (Duchêne et al., 2015, Rieux and Balloux, 2016), although the presence and direction of a potential bias remain poorly understood. However, it is important to note that a lack of temporal signal does not necessarily preclude estimating evolutionary rates and timescales because alternative sources of calibration, such as prior estimates of evolutionary rates or constraints on internal node ages, can still be used to inform analyses.

In principle, frameworks developed to test for temporal signal do not differentiate between recently emerging organisms (Fig.1a) and those with narrow sampling windows (Fig.1d), both of which may lack temporal signal. As most of these tests involve fitting a phylogenetic model to the data, they implicitly assume that the model adequately captures the evolutionary process and thus their performance also highly depends on model fit. Recent research, for example, suggests that the choice of tree prior and molecular clock model substantially impacts the sensitivity and specificity of temporal signal tests (Tay et al., 2024). Thus, temporal signal is not solely a property of the data but also depends on the choice of model.

**Figure 1.**
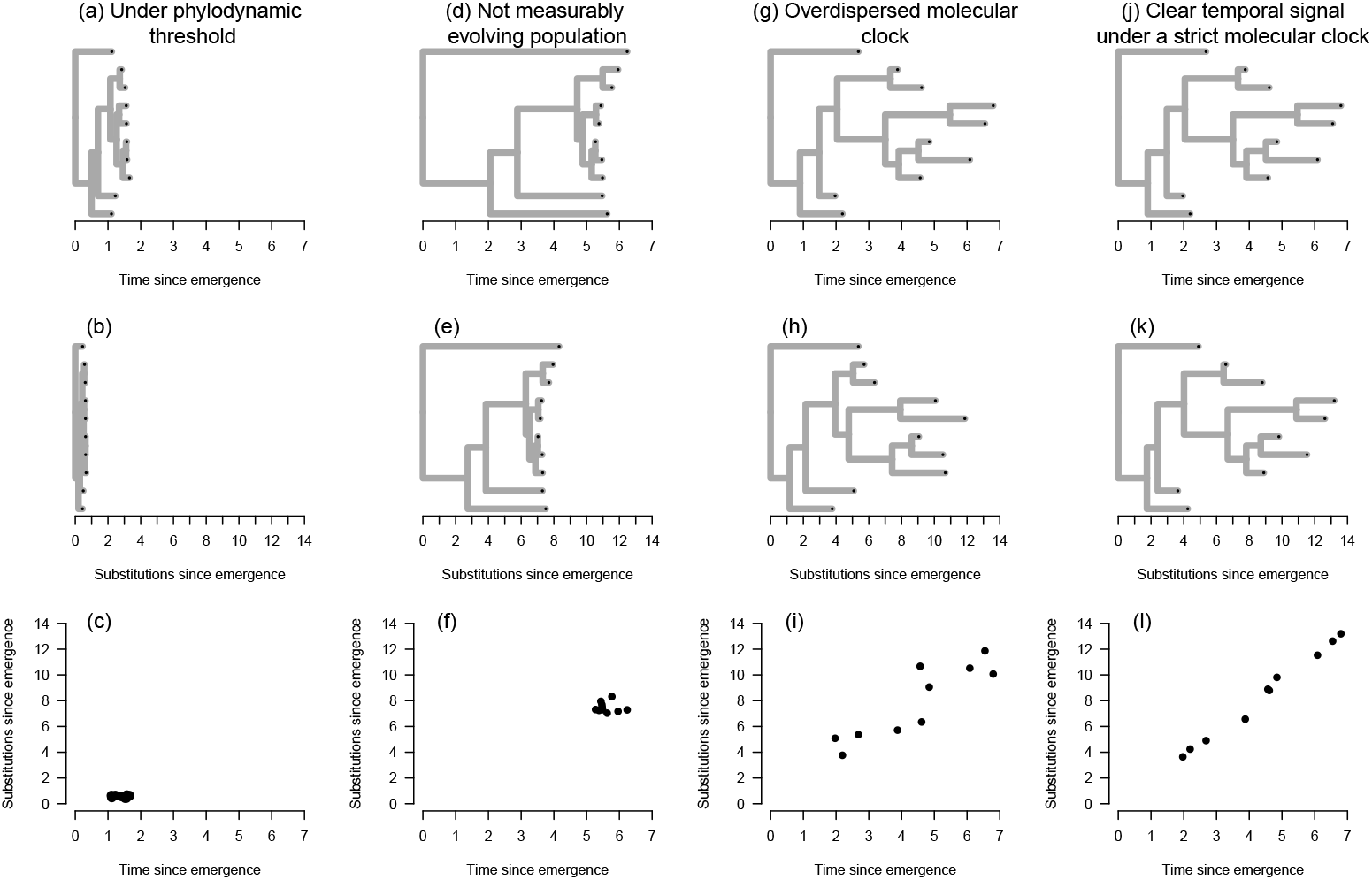
Examples of situations where temporal signal may or may not be detected. An organism that has not attained its phylodynamic threshold has a recent time of emergence (with a phylogenetic time-tree shown in (a)) because it has not had sufficient time to accrue an appreciable number of substitutions (phylogenetic tree with branch lengths in subs/site, i.e. a ‘phylogram’, shown in (b)), such that it is not possible to establish a statistical relationship between molecular evolution (substitutions) and time (shown in (c)). Sequence data from an organism that has evolved for a substantial amount of time may have been sampled over a very narrow window of time that is not sufficient to treat it as a measurably evolving population (time-tree in (d) and phylogram in (e)), which results in no temporal signal (root-to-tip regression in (f)). A data set may involve a wide sampling window of time and from a population that has attained its phylodynamic threshold, but an overdispersed molecular clock (substantial rate variation among lineages; panels (g) - (i)) may result in a lack of temporal signal. In (j) through (l) we show the situation where an organism has attained its phylodynamic threshold, it has been sampled for sufficiently long, and where evolutionary rate variation among lineages is negligible, as to produce a clear relationship between molecular evolution and time, and thus unequivocal temporal signal.

Various methods exist for assessing temporal signal. The root-to-tip regression (Buonagurio et al., 1986, Drummond et al., 2003a, Gojobori et al., 1990) fits a regression to the distance from the root to the tips in a phylogenetic tree against sampling time. High R2 values of the regression suggest that phylogenetic distance can be sensibly modelled as a linear function of time and can therefore be used as an indication of informative tip-calibrations. Date-randomisation tests (Duchêne et al., 2015, Duchene et al., 2018, Ramsden et al., 2009, Trovão et al., 2015) compare evolutionary rate estimates using correct sampling times against those from permutations. Bayesian Evaluation of Temporal Signal (BETS; Duchene et al. (2020c)) evaluates whether a model with sampling times performs better than a model that assumes isochronous sampling using Bayes factors. Each method comes with a set of limitations and strengths, such that tests of temporal signal should be used in combination, rather than being mutually exclusive (Duchene et al., 2020c, Rieux and Balloux, 2016).

### 1.4 Concepts of measurably evolving populations, the phylodynamic threshold, and temporal signal in practice

In Fig.1, we present four simple example cases to illustrate the relationships among the concepts of mea surably evolving populations, the phylodynamic threshold, and temporal signal. The first example depicts an organism that has emerged recently and therefore has not yet reached its phylodynamic threshold (with a phylogenetic time-tree shown in panel (a)). Due to its recent origin, there has not been enough time for the accumulation of a sufficient number of substitutions (represented in the phylogram in panel (b)), and thus it is not possible to establish a statistical relationship between molecular evolution (i.e., substitutions) and time (as shown in panel (c)). A real-world example of such a case comes from the early phase of the SARS-CoV-2 outbreak: initial efforts to estimate the evolutionary rate and time of origin had substantial uncertainty due to a narrow sampling window and low genetic diversity (Boni et al., 2020). In Duchene et al. (2020a), Bayesian phylodynamic analyses were conducted on genome data as the outbreak unfolded. The number of available genomes and the width of the sampling window increased over time and ranged from 22 genomes sampled over 31 days to 122 genomes sampled over 63 days. Although early estimates of the evolutionary rate and time of origin were highly uncertain, they quickly converged to stable values as more data became available (Ghafari et al., 2022). Duchene et al.

The second example in Fig.1 illustrates a case in which an organism has evolved over a long period, but the available sequence data have been collected within a very narrow window of time, insufficient to treat the data set as a measurably evolving population (time-tree in panel (d) and phylogram in panel (e)). This results in no temporal signal, as demonstrated by the lack of correlation in the root-to-tip regression in panel (f). The causative agent of tuberculosis, the bacterium *Mycobacterium tuberculosis*, was commonly considered to evolve too slowly for calibrating the molecular clock using samples collected over a few years (2016). A range of studies have shown, however, that for *M. tuberculosis* a genome sampling window of a few decades might be sufficient for reliable clock calibration (Eldholm et al., 2015, Kühnert et al., 2018, Menardo et al., 2019, Merker et al., 2022).

The third example in Fig.1 describes a data set that may involve a wide sampling window of time and for which samples have been drawn from a population that has attained its phylodynamic threshold, but with substantial rate variation among lineages – i.e. overdispersed molecular clock –, resulting in a lack of temporal signal (panels (g) – (i)). This pattern appears to be the case in *Yersinia pestis*, the bacterium that causes the plague, for which some localised outbreaks display obvious temporal signal, but its long-term evolution has pervasive evolutionary rate variation (Andrades Valtuenã et al., 2022, Eaton et al., 2023).

In the final example in Fig.1, a hypothetical organism has attained its phylodynamic threshold, has been sampled for sufficiently long time, and evolutionary rate variation among lineages is low. These conditions together produce a clear relationship between molecular evolution and time, thus providing unequivocal temporal signal (panels (j) – (l)). The long term evolution of *Vibrio cholerae*, the causative agent of cholera, and H3N2 influenza virus are exemplar microbes whose molecular evolution has been fairly constant across long periods of time (Devault et al., 2014, Rambaut et al., 2016).

In summary, the concepts of measurably evolving population, phylodynamic threshold and temporal signal describe the information that can be drawn from a sampled population about its evolutionary timescale. Because populations that are not measurably evolving have been observed to yield biased estimates, they remain important to consider (Gharbi et al., 2024). In Bayesian inference, such biases can be the result of complex interactions between prior distributions or model settings that do not align with the true data generating model, as these drive the inference in the absence of informative data. Traditionally, potential biases due to prior interactions (Tay et al., 2024) or model misspecification (Möller et al., 2018) have been found through simulation studies, while data analyses often involve little validation of the results (Mendes et al., 2025). Here, we illustrate through a range of examples the degree to which differing levels of temporal signal in a data set can interact with prior settings and model assumptions, both on simulated and empirical data.

## 2 Results

We sought to pinpoint the impact of sampling strategies on molecular clock estimates. We focused our attention on two major problems for emerging microbes and studies involving ancient DNA. First, we conducted simulations varying the sampling window of a population that had attained its phylodynamic threshold, and which was analysed with a range of prior distributions on the evolutionary rate. In the second simulation scenario, we subsampled a population over time to vary the number of ancient samples, leading to a temporal sampling bias. Finally, we illustrate these results in an empirical data set of Hepatitis B virus (HBV) that includes a large number of ancient samples (Kocher et al., 2021). This virus has been the subject of intense research due to its close association with human populations and complex evolutionary dynamics (Kahila Bar-Gal et al., 2012, Paraskevis et al., 2013, Ross et al., 2018).

### 2.1 Sampling windows relative to the phylodynamic threshold

We simulated sequence data that resembled the evolution of HBV, a double-stranded DNA (dsDNA) virus that has evolved in humans for at least ten thousand years (Kocher et al., 2021). Our synthetic data had a genome length of 3,200 nucleotides and an evolutionary rate of 1.5 × 10^−5^ subs/site/year (Kocher et al., 2021, Mühlemann et al., 2018) with a moderate amount of rate variation among lineages (see Materials and methods). Under these conditions, we expect to observe one mutation every 20 years 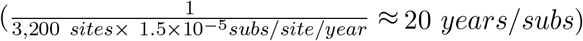. This number is important for the design of our simulation experiments: 20 years is the expected phylodynamic threshold, as introduced above, and typically serves as a good reference point from when to expect temporal signal. We analysed the data under a Bayesian phylogenetic framework and computed three performance statistics from the posterior: coverage (whether the 95% credible interval, CI, contains the true value used to generate the data), uncertainty (the width of the posterior 95% CI divided by its mean), and bias (the difference between the posterior mean and the truth divided by the truth).

We conceived a simulation process under which the evolutionary timescale had an expectation of ten thousand years and with a sampling window of 0, 10, 20, 200, or 2,000 years. A sampling window spanning 0 years results in ultrametric trees with the sampling times providing no calibration information (all calibration information comes from the tree prior and the prior on the evolutionary rate). In contrast, a sampling window of 10 years is half of the expected phylodynamic threshold and is likely to have weak temporal signal (see Fig.1(d)-(f)). Sampling windows of 20 years (the expected phylodynamic threshold) or wider are more likely to behave as measurably evolving populations with increasingly strong temporal signal (see Fig.1(j)-(l)). Our synthetic data sets were analysed under a Bayesian phylogenetic framework, as implemented in the BEAST2 platform (Bouckaert et al., 2019).

To investigate the impact of the prior we considered several configurations for the prior on the mean evolutionary rate. In our analyses the molecular clock model is an uncorrelated relaxed molecular clock model with an underlying lognormal distribution, with mean *M*. For this parameter we set nine possible prior Gamma distributions, for which the prior mean could be the value used to generate the data, or one order of magnitude higher or lower. We also included three degrees of uncertainty in this prior (see Fig.2 and table 6). In this respect, a prior with low uncertainty, and a mean that is much higher or lower than the true should result in more bias than one that has higher uncertainty or is centred on the true value.

Our analyses for which the prior on *M* was centred on the true value had very high coverage. For each sampling window, the analysis recovered the true value of *M* within the 95% CI in at least 94 out of 100 simulated datasets (table and Fig. 2). However, when the prior was highly precise (95% CI/mean=1.0) but biased downwards, even the simulations with a sampling window of 100× the phylodynamic threshold (i.e. 2,000 years before present) still yielded low coverage (only 1 out of 100 analyses of the simulation replicates included the true value in the 95% credible interval, table 1).

**Table 1:** Coverage of the mean evolutionary rate, *M*. We show the number of simulation replicates out of 100 for which we found coverage. Data were simulated under trees with five possible sampling window depths relative to the expected phylodynamic threshold, where Sw=*D*× is for simulations with a sampling window of *D* times the expected phylodynamic threshold (20 years in our data; see Fig. 2). Each column is a sampling window depth, and the rows denote the configuration of the prior on the mean rate, *M*, for which we show the prior mean and the width of the 95% CI divided by the mean (larger values imply higher uncertainty). The true value of *M* is 1.5 × 10^−5^ subs/site/year. The horizontal lines separate prior configurations with different prior means.

| Prior configuration<br>(mean, 95%CI width) | Sw=0× | Sw=0.5× | Sw=1× | Sw=10× | Sw=100× |
| --- | --- | --- | --- | --- | --- |
| $1.5 \times 10^{-5}$, 1.00 | 100 | 100 | 100 | 94 | 94 |
| $1.5 \times 10^{-5}$, 3.04 | 100 | 100 | 100 | 96 | 95 |
| $1.5 \times 10^{-5}$, 6.33 | 100 | 100 | 99 | 95 | 94 |
| $1.5 \times 10^{-4}$, 1.00 | 0 | 0 | 0 | 77 | 91 |
| $1.5 \times 10^{-4}$, 3.04 | 100 | 97 | 97 | 93 | 93 |
| $1.5 \times 10^{-4}$, 6.33 | 100 | 100 | 99 | 94 | 92 |
| $1.5 \times 10^{-6}$, 1.00 | 0 | 0 | 0 | 0 | 1 |
| $1.5 \times 10^{-6}$, 3.04 | 0 | 0 | 0 | 68 | 91 |
| $1.5 \times 10^{-6}$, 6.33 | 100 | 88 | 85 | 95 | 94 |

**Figure 2.**
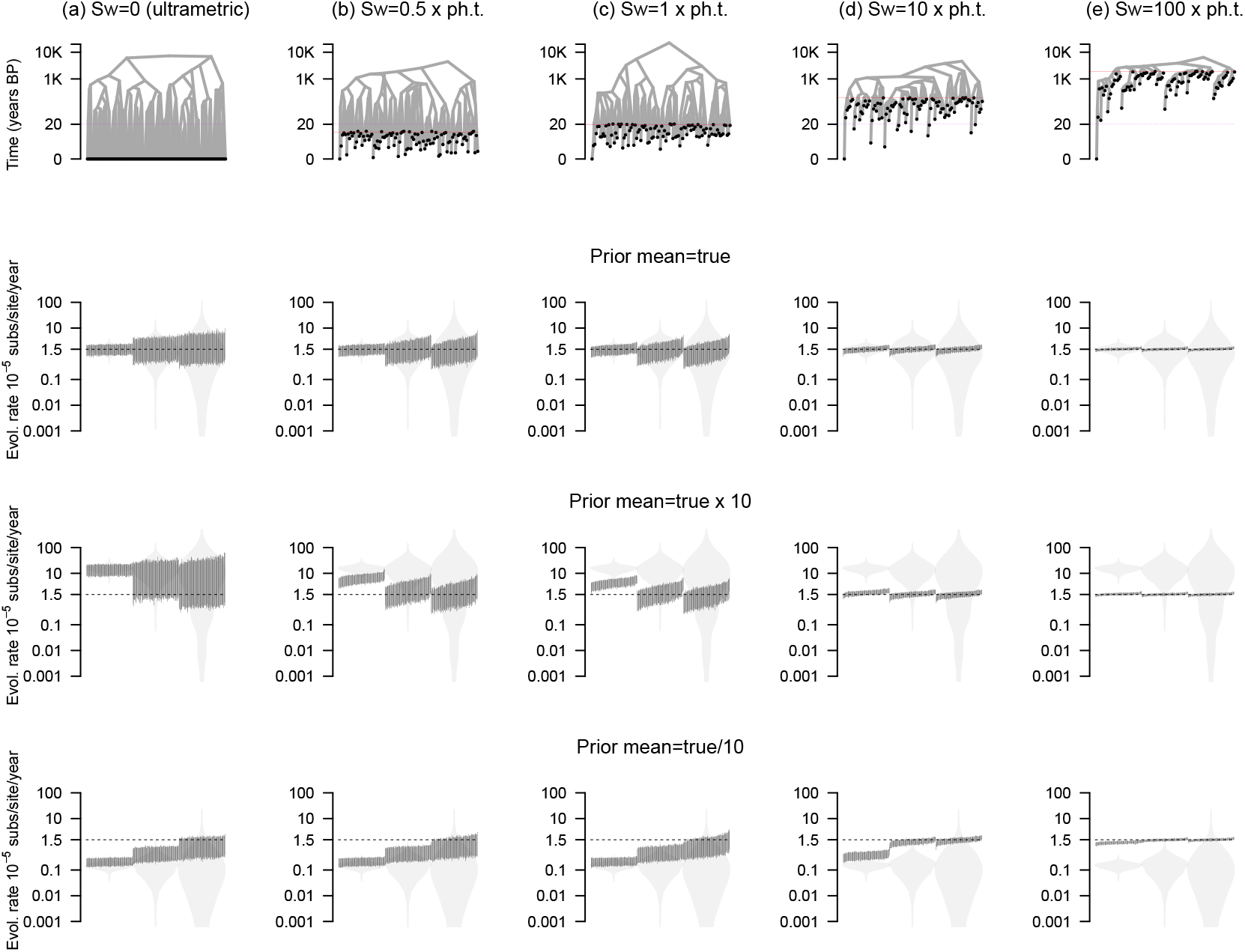
Estimates of evolutionary rates, *M* for simulations of varying sampling window widths. Each column corresponds to a simulation setting: (a) is for ultrametric trees where all samples are collected at the same point in time (sampling window, sw=0), (b) is for the situation where the sampling window is 10 years (half the expected phylodynamic threshold; sampling window, sw=0.5× ph.t), (c) is where the sampling window is exactly the expected phylodynamic threshold of 20 years (sw=1× ph.t). Scenarios (d) and (e) denote sampling windows that are 10 and 100 × the expected phylodynamic threshold (sw=10× ph.t and sw=100× ph.t, respectively). The rows denote example phylogenetic trees and prior configurations where the mean is set to the correct value (first row, 1.5 × 10^−5^), an order of magnitude higher (second row, 1.5 × 10^−4^), or an order of magnitude lower (last row, 1.5 × 10^−6^). The prior distributions are shown with the grey violins (from left to right in each plot, uncertainties are 1.00, 3.04, and 6.33) and each black bar is the 95% credible interval of the posterior. The dashed line in each case denotes the correct value.

The uncertainty in estimates of evolutionary rates was associated with the width of the sampling window, but also with the uncertainty in the prior. In the situation where the sampling window was 1× the expected phylodynamic threshold or less we consistently found that uncertainty in the prior was commensurate with uncertainty in the posterior. Estimates using a prior uncertainty of 1.00 (the width of the 95% CI is the same as the mean) are narrower than those using a prior of uncertainty of 3.04 or 6.33 (see table 2 and Fig.).

**Table 2:** Average uncertainty of the mean evolutionary rate, *M* across 100 simulation replicates in each case. We quantify uncertainty as the posterior width of the 95% credible interval (CI) divided by the mean. The rows and columns match those from table 1, with the posterior for each simulation replicate shown in Fig. 2.

| Prior configuration<br>(mean, 95%CI width) | Sw=0× | Sw=0.5× | Sw=1× | Sw=10× | Sw=100× |
| --- | --- | --- | --- | --- | --- |
| $1.5 \times 10^{-5}$, 1.00 | 0.96 | 0.94 | 0.91 | 0.45 | 0.21 |
| $1.5 \times 10^{-5}$, 3.04 | 2.24 | 1.87 | 1.55 | 0.48 | 0.21 |
| $1.5 \times 10^{-5}$, 6.33 | 3.06 | 2.01 | 1.72 | 0.48 | 0.22 |
| $1.5 \times 10^{-4}$, 1.00 | 1.04 | 0.87 | 0.80 | 0.43 | 0.21 |
| $1.5 \times 10^{-4}$, 3.04 | 1.36 | 1.83 | 1.50 | 0.48 | 0.22 |
| $1.5 \times 10^{-4}$, 6.33 | 5.39 | 2.11 | 1.70 | 0.49 | 0.21 |
| $1.5 \times 10^{-6}$, 1.00 | 0.75 | 0.75 | 0.76 | 0.75 | 0.28 |
| $1.5 \times 10^{-6}$, 3.04 | 1.36 | 1.37 | 1.38 | 0.52 | 0.22 |
| $1.5 \times 10^{-6}$, 6.33 | 1.99 | 1.90 | 1.72 | 0.48 | 0.22 |

When the sampling window was 10× the phylodynamic threshold or more we found a more complicated picture. When the prior had a downward bias (mean=1.5 × 10−6 subs/site/year) and a low uncertainty of 1.00, the posterior estimate was wider than when this prior was less uncertain (table 2 and Fig. 2). This finding can be explained because a very wide sampling window provides a large amount of information for inferring the evolutionary rate, which can yield high uncertainty if the prior stands in conflict. Importantly, the widest sampling window in our experiments, of 100× the expected phylodynamic threshold produced consistently low uncertainty, although it is important to note that when the prior is highly biased downward (mean of 1.5 × 10−6 subs/site/year and uncertainty of 1.00) coverage is very low, with only one simulation replicate containing the true value used to generate the data within the 95% CI.

To understand the directionality of posterior evolutionary rate estimates relative to the value used to generate the data, we quantified the amount of bias, as the difference between the true evolutionary rate and the posterior mean divided by the true value (i.e. 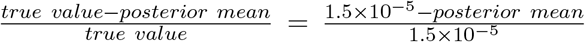. As expected, when the prior was centred in the true value, we observed minimum bias. Across all prior configurations, average bias values ranged from 0.00 to 8.31 (a posterior estimate that was on average up to 8 times higher than the truth). The most marked average bias was found for our priors with an uncertainty of 1.00 and a mean of 1.5×10^−4^*subs/site/year* and 1.5×10^−6^*subs/site/year*, respectively (table 3). Increasingly wide sampling windows had lower amounts of bias. A sampling window of 100× the phylodynamic threshold had a maximum average bias of -0.22 for the prior with downward bias and low uncertainty.

**Table 3:** Average bias of the mean evolutionary rate, *M* across 100 simulation replicates in each case. Bias is calculated as the true evolutionary rate, 1.5 × 10^−5^ subs/site/year - posterior mean, divided by the true value.

| Prior configuration<br>(mean, 95%CI width) | Sw=0× | Sw=0.5× | Sw=1× | Sw=10× | Sw=100× |
| --- | --- | --- | --- | --- | --- |
| $1.5 \times 10^{-5}$, 1.00 | 0.00 | -0.01 | -0.01 | 0.00 | 0.01 |
| $1.5 \times 10^{-5}$, 3.04 | 0.16 | 0.06 | 0.01 | 0.00 | 0.01 |
| $1.5 \times 10^{-5}$, 6.33 | 0.28 | 0.03 | -0.03 | 0.00 | 0.01 |
| $1.5 \times 10^{-4}$, 1.00 | 8.31 | 3.22 | 1.84 | 0.17 | 0.04 |
| $1.5 \times 10^{-4}$, 3.04 | 4.93 | 0.67 | 0.30 | 0.02 | 0.01 |
| $1.5 \times 10^{-4}$, 6.33 | 3.37 | 0.27 | 0.08 | 0.00 | 0.01 |
| $1.5 \times 10^{-6}$, 1.00 | -0.86 | -0.86 | -0.86 | -0.75 | -0.22 |
| $1.5 \times 10^{-6}$, 3.04 | -0.71 | -0.70 | -0.69 | -0.16 | -0.02 |
| $1.5 \times 10^{-6}$, 6.33 | -0.45 | -0.43 | -0.37 | -0.04 | 0.00 |

Overall, these simulations demonstrate that increasingly wide sampling windows result in evolutionary rate estimates that are more accurate, precise, and less biased, than those from data sets with narrow sampling windows. Although increasingly wide sampling windows are more robust to prior misspecification, we emphasise the importance of choosing the prior for the evolutionary rate carefully because a highly biased prior can overcome the information from data with wide sampling windows. Contrary to the expectation that low temporal signal necessarily results in an underestimation of the evolutionary rate and an overestimation of the tree height (Duchêne et al., 2015), we find that a lack of temporal signal due to narrow sampling windows may simply lend more influence to the prior.

For all trees that were non-ultrametric (i.e. those with a sampling window width of at least 0.5× the phylodynamic threshold) a downward bias in the evolutionary rate prior appears to be more detrimental than one with an upward bias. In our simulations with sampling windows of width 100× the phylodynamic threshold, a highly downwards biased prior (mean=1.5 × 10^−6^ subs/site/year and uncertainty of 1.00) still had very low coverage. In contrast, a prior with a similar upward bias (mean=1.5 × 10^−4^ and uncertainty of 1.00) produced much higher coverage for sampling windows from 10× the phylodynamic threshold (table 1).

These results indicate that priors with high uncertainty should be advised for practical studies where a researcher is unsure about the evolutionary rate of the population. In our simulations posterior estimates using a prior uncertainty of 6.33 seemed to produce a good trade-off between uncertainty and accuracy. Such a prior means that the 95% CI spans just over six orders of magnitude. For a sampling window of 1× the expected phylodynamic threshold, the posterior distribution had an average uncertainty of around 1.7, which may be sufficient for biological interpretation of estimates of evolutionary rates and timescales, under these particular simulation settings.

For completeness, we also analysed our simulated data with a uniform prior on *M* bounded between zero and infinity. This is a default prior in some versions of BEAST2, but we note that it is unadvisable for several reasons. First, it is not a proper probability distribution because the area under the curve is not one, such that it can lead to unreliable model selection using marginal likelihoods and Bayes factors (Fourment et al., 2020, Oaks et al., 2019). Second, although this distribution is commonly perceived to be ‘uninformative’, there is an infinite amount of density on implausible high values. In our analyses we found that for the ultrametric trees the Markov chain often had unreliable mixing (low effective sample size), with mean posterior estimates for *M* that could be as high as 1 × 10^50^ subs/site/year. However, those simulations with a sampling window of 0.5 × the phylodynamic threshold produced estimates that were much more reasonable, with subsequently wider sampling windows producing estimates with lower uncertainty and higher accuracy (see Supplementary material online).

### 2.2 Hierarchical priors and the phylodynamic threshold

An attractive approach for specifying prior distributions for parameters that are largely unknown is to use a hierarchical structure, where the parameters that govern the prior have priors themselves (sometimes known as hyperpriors). In the context of Bayesian molecular clocks, such hierarchical structure has been used to specify uncertainty in time calibrations (Heath et al., 2014). We employed this method by setting a prior for the shape and rate parameters of the Gamma distribution that governs the evolutionary rate, *M* (see Materials and methods).

Under the hierarchical structure, the marginal prior for *M* is very uncertain and is biased upwards, relative to the true value (Fig.3). However, for sampling windows of at least 1× the phylodynamic threshold its performance in terms of coverage, average uncertainty, and average bias were comparable to those using the standard prior centred on the correct value (see first three rows under SW= 1× in tables 1, 2, and 3, compared to table 4). This result demonstrates the effect of Bayesian regularisation, where the model is effectively able to learn the parameters of the prior of the evolutionary rate, and that using hierarchical models is likely preferable to using a prior that is potentially highly biased.

**Figure 3.**
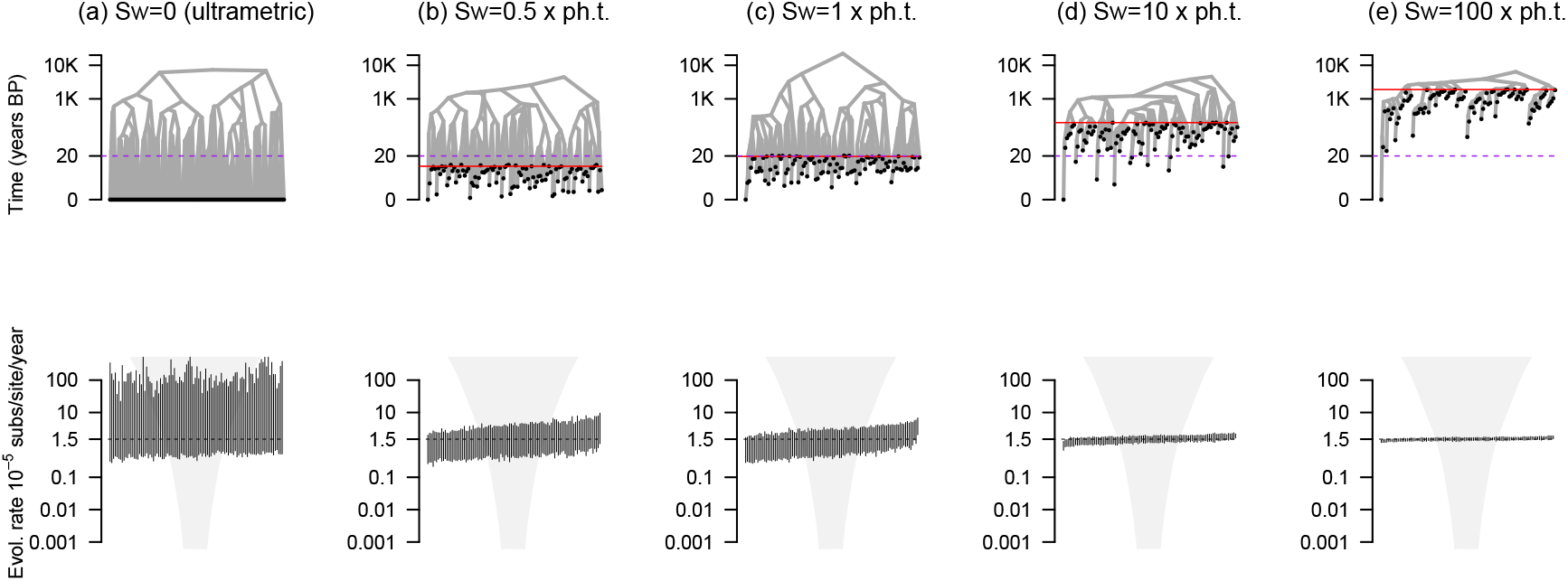
Estimates of evolutionary rates, *M* for simulations of varying sampling window widths. Each column corresponds to a simulation setting: (a) is for ultrametric trees where all samples are collected at the same point in time (sampling window, sw=0), (b) is for the situation where the sampling window is 10 years (half the expected phylodynamic threshold; sampling window, sw=0.5× ph.t), (c) is where the sampling window is exactly the expected phylodynamic threshold of 20 years (sw=1× ph.t). Scenarios (d) and (e) denote sampling windows that are 10 and 100 × the expected phylodynamic threshold (sw=10× ph.t and sw=100× ph.t, respectively). The first row denotes example phylogenetic trees, while the second row is the corresponding posterior estimates. The prior is shown with the grey violins (a hierarchical prior configuration) and each black bar is the 95% credible interval of the posterior. The dashed line in each case denotes the correct value.

**Table 4:**
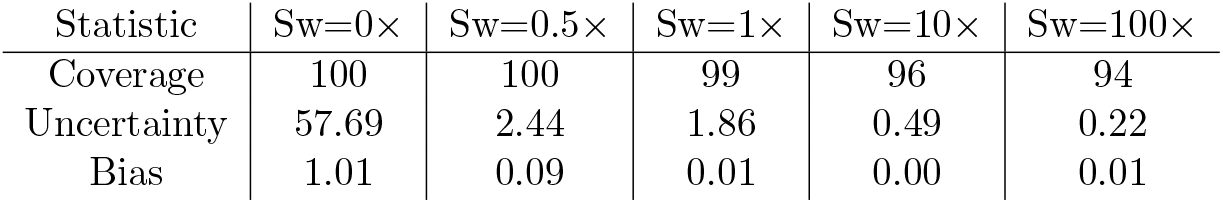
Coverage, average uncertainty and average bias of the mean evolutionary rate, *M* using a hierarchical prior on *M*. Statistics are reported over 100 simulation replicates under each of the five sampling window widths (Sw) relative to the phylodynamic threshold (of 20 years).

| Statistic | Sw=0× | Sw=0.5× | Sw=1× | Sw=10× | Sw=100× |
| --- | --- | --- | --- | --- | --- |
| Coverage | 100 | 100 | 99 | 96 | 94 |
| Uncertainty | 57.69 | 2.44 | 1.86 | 0.49 | 0.22 |
| Bias | 1.01 | 0.09 | 0.01 | 0.00 | 0.01 |

### 2.3 Temporal sampling bias

We investigated the impact of temporal sampling bias on the quality of molecular clock estimates. For this purpose we simulated data with the same genomic characteristics as HBV and where genome sampling was conducted over five periods of time uniformly distributed between the present and the root of the tree (Fig.4(a)). The fully sampled trees contained 500 genome samples, with 100 for each of the five sampling times. Such stratified sampling is expected in ancient DNA studies, for example when a set of samples are drawn from archaeological sites (e.g. Spyrou et al. 2019b).. We sampled the complete data sets either by randomly selecting 20 samples from each strata, which we refer to as ‘time-uniform’ sampling, or by sampling with a probability that is inversely proportional to the age of the strata, referred to as ‘time-biased’. The time-uniform and time-biased sampling strategies both contain 100 samples (1/5^*th*^ of the complete data), but the time-biased only includes a small number of ancient samples.

**Figure 4.**
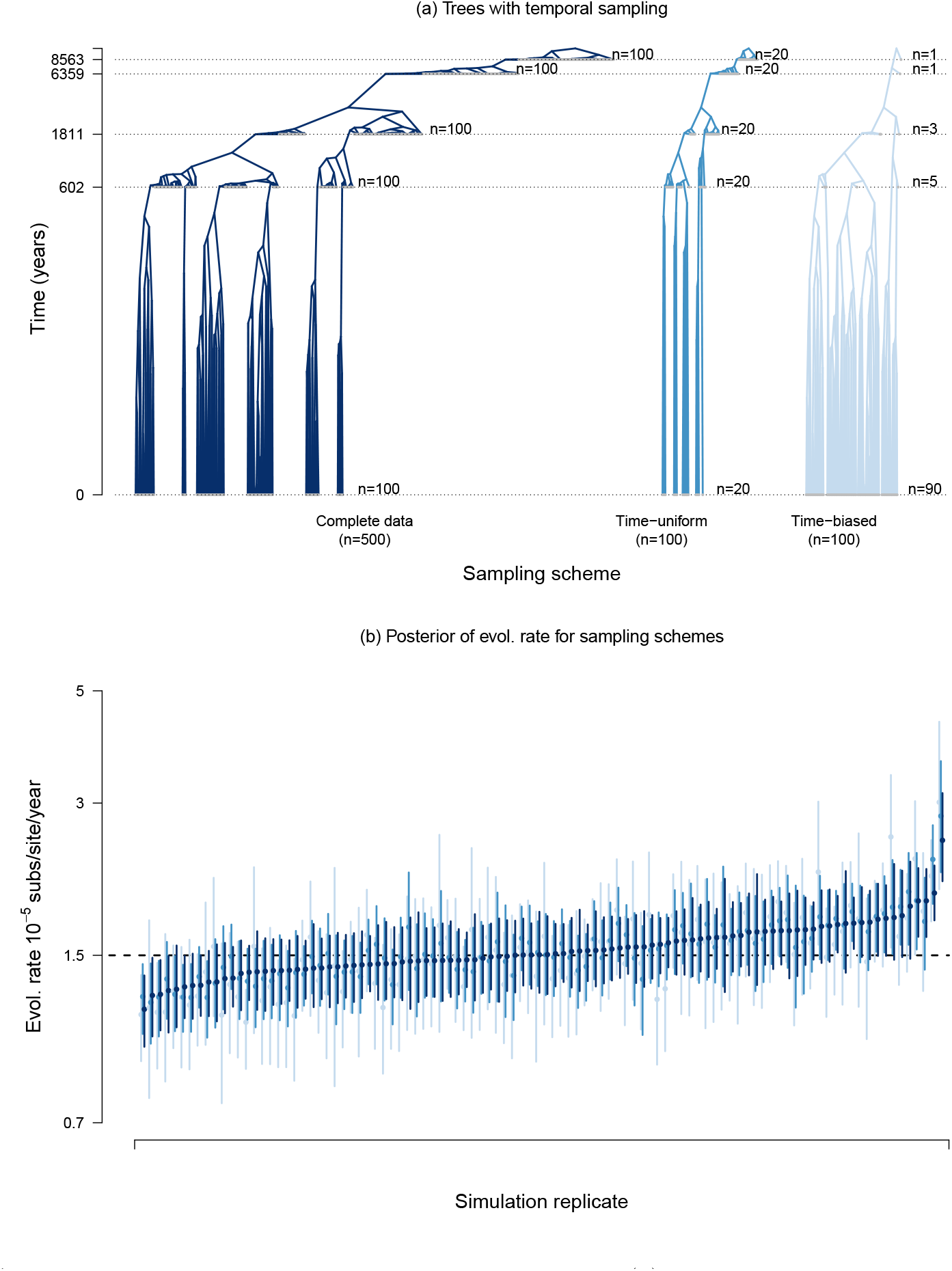
Analyses under sampling treatments over time. In (a) we show an example of the trees for a simulation replicate, with branch lengths and time in log_10_ scale. The complete data set consists of 500 genome samples, collected in five points in time, with an equal number of samples per time point (n=100). The first sampling strategy is unbiased, where 20 samples are drawn from each time point, and is known here as ‘time-uniform’. The ‘time-biased’ regime is where sampling intensity decreases over time. Note that the total number of samples in the time-uniform and time-biased treatments is identical. In (b) we show the posterior estimates of the evolutionary rate for each treatment. Each simulation replicate is represented by three error bars: dark blue for the complete data, and lighter shades of blue for the estimates from the time-uniform and time-biased sampling treatments. The width of the error bars denotes the 95% quantile range and the dots are the mean value. The dashed line shows the true value used to generate the data.

The coverage of the evolutionary rate estimate was comparable across simulation treatments (table 5). It is worth noting that the coverage and other performance metrics were lower than for our simulation treatment where samples are uniformly drawn between the phylodynamic threshold and the present, which likely occurs because in this case samples with identical sampling times form monophyletic groups, reducing the amount of temporal information in the data (see Murray et al. 2016 for a detailed investigation). Our measures of coverage and average bias did not differ substantially between sampling treatments.

**Table 5:**
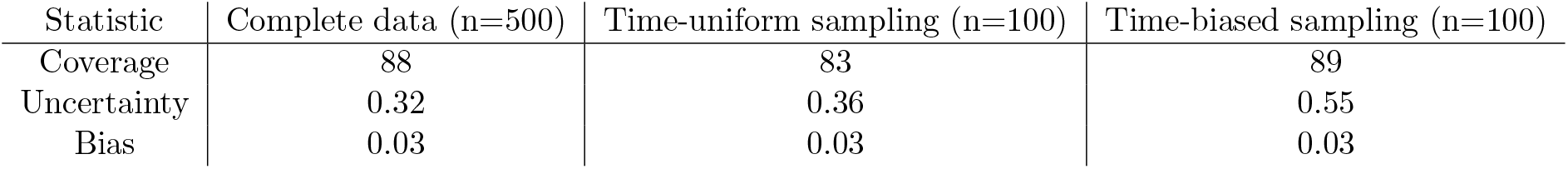
Coverage, average uncertainty and average bias of the mean evolutionary rate, *M*, for data with temporal sampling bias. Statistics are reported over 100 simulation replicates for three possible scenarios, shown in Fig.4(a).

| Statistic | Complete data (n=500) | Time-uniform sampling (n=100) | Time-biased sampling (n=100) |
| --- | --- | --- | --- |
| Coverage | 88 | 83 | 89 |
| Uncertainty | 0.32 | 0.36 | 0.55 |
| Bias | 0.03 | 0.03 | 0.03 |

A striking result of the temporal sampling strategies was its impact on the uncertainty of the posterior. Both the time-uniform and time-biased sampling treatments resulted in posterior distributions that were wider than with the complete data, which is to be expected because they are effectively smaller data sets with less information, but with no systematic bias (Fig.5(a)). However, the time-biased sampling data sets almost invariably have posterior distributions with higher uncertainty than those from the time-uniform sampling (Fig.5(b)), implying that the distribution of samples, and not just the number, is important for estimation uncertainty. We note that in empirical analyses of ancient DNA data systematic biases may occur with the inclusion of ancient samples, particularly due to sequencing errors that are more likely in ancient samples, and therefore a careful assessment of their impact is advisable, for example by conducting analyses where these samples are sequentially added.

**Figure 5.**
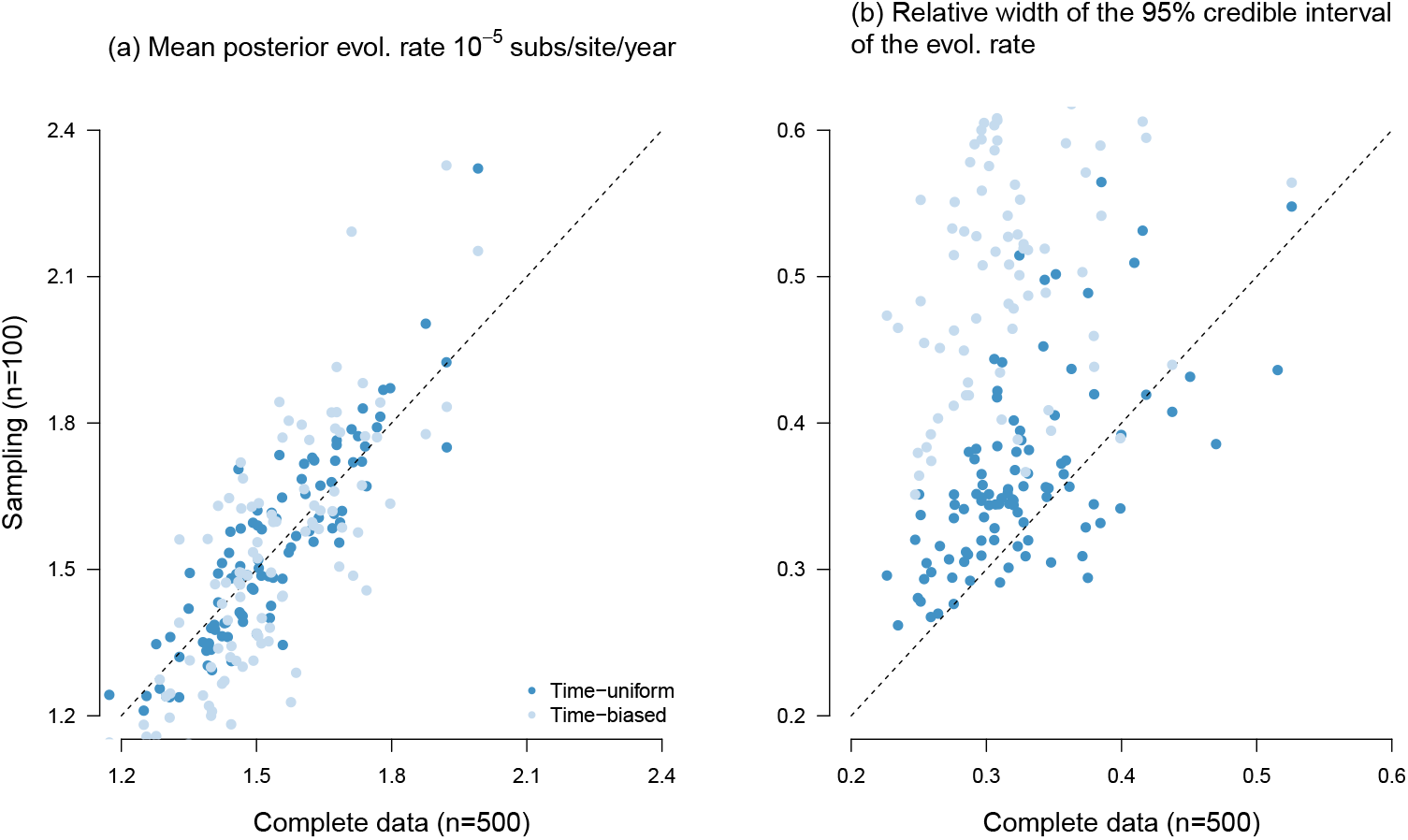
Comparison of posterior evolutionary rate estimates between complete data (x-axis) and two sampling treatments (y-axis): time-uniform (dark blue) and time-biased (light blue). Each dot is a simulation replicate. In (a) we show the mean posterior evolutionary rate, *M*, estimate. Points that fall along the y=x line (dashed line) represent identical mean posterior for the sampling treatment and the complete data, while those above or below represent higher or lower estimates, respectively, relative to the complete data. In (b) we show the uncertainty, calculated as the width of the lower 95% credible interval divided by the mean value. Values that fall along the y=x line denote those for which the complete data and either sampling strategies are equally uncertain, while those above and below the y=x line are more or less uncertain, respectively.

### 2.4 Empirical analyses of Hepatitis B virus (HBV) ancient and modern genomes

To explore the impact of the width of the sampling window and the temporal sampling bias on the estimates of evolutionary rates and times, we performed analyses of a HBV data set that includes modern and ancient genomes, from Kocher et al. (2021). The complete data set consisted of 232 genomes of length 3,344 nucleotides and with a sampling window of 10,535 years. HBV is an ancient pathogen that has likely codiverged with human populations for thousands of years (Locarnini et al., 2021, Mühlemann et al., 2018, Paraskevis et al., 2013, Zehender et al., 2014), and thus its phylodynamic threshold has been reached, although it has not been empirically established whether it can be considered to be a measurably evolving population, as has also been the case for recent outbreaks like SARS-CoV-2 (Duchene et al., 2020a).

We analysed the data by drawing 100 sequences with replacement and restricting their sampling window. We constructed six of such data sets, with sampling windows of 10 years (approximately half of the expected phylodynamic threshold), 19 years (approximately the expected phylodynamic threshold), 32, 422, 1,863 and 10,526 years. Increasing the sampling window resulted in estimates of the evolutionary rate that were more precise and closer to the estimate from the complete data set (Fig.6). Here we find that the evolutionary rate is estimated to be higher for shorter sampling windows, with correspondingly younger estimates for the tree height. This pattern can be due to one or a combination of other factors influencing the inference, for example, the vagaries of evolutionary rate variation in this virus, particularly time-dependency (Vrancken et al., 2017). Similarly, population structure that is unaccounted for has been shown to produce an overestimation of the evolutionary rate, because under the tree prior samples that are genetically linked are expected to have been sampled at the same point in time (Möller et al., 2018).

**Figure 6.**
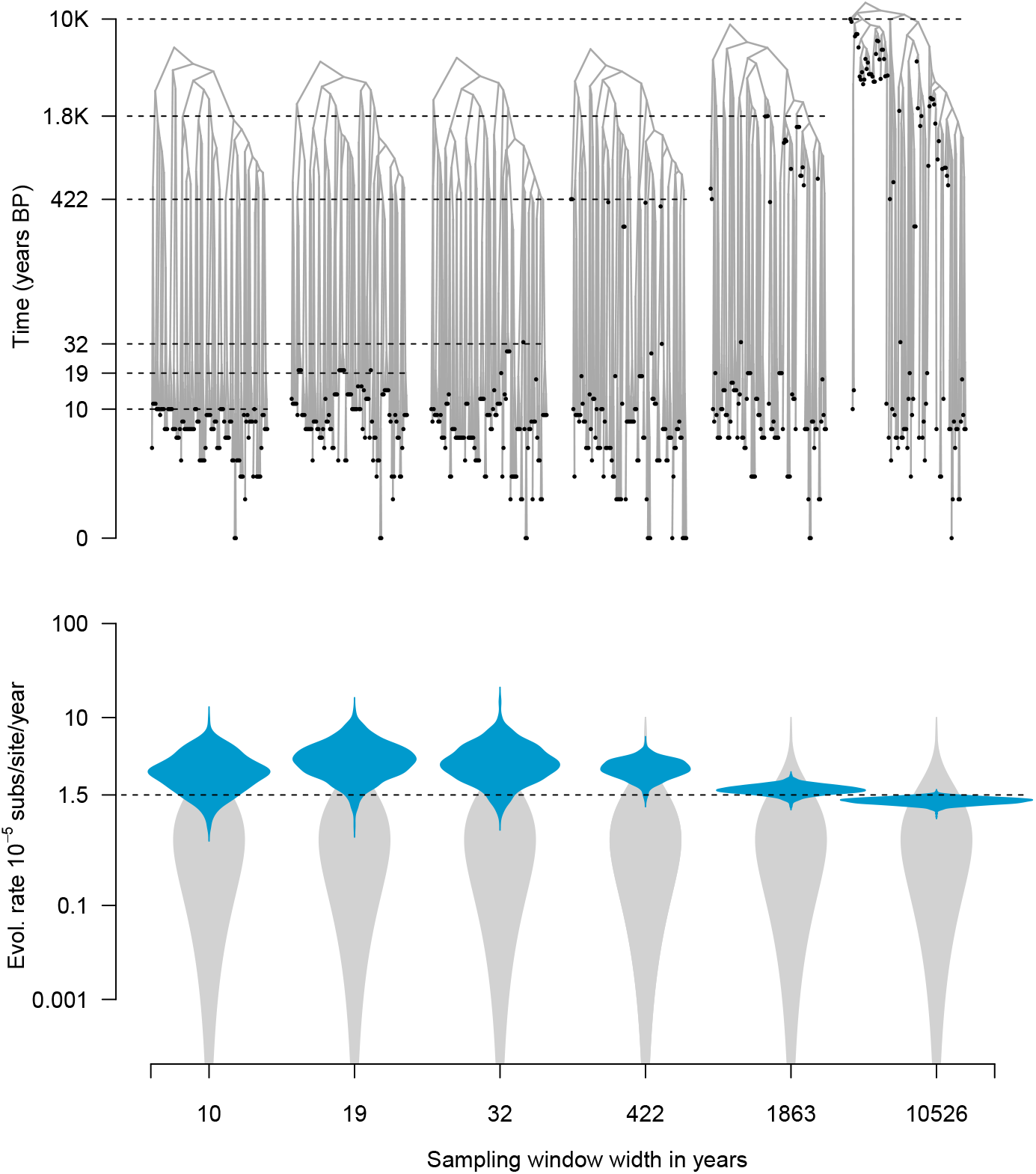
Results from empirical analyses of Hepatitis B virus (HBV) ancient DNA data. The phylogenetic trees correspond to highest clade credibility trees from six analyses with sampling windows of 10 years (approximately half of the expected phylodynamic threshold), 19 years (approximately the expected phylodynamic threshold), 32, 422, 1,863 and 10,526 years. The violin plots show the posterior distribution of the evolutionary rate, *M*, in blue and its corresponding prior in grey. The dashed line shows the mean *M* estimate from the complete data set.

## 3 Discussion

The concepts of the phylodynamic threshold, measurably evolving populations, and temporal signal are helpful for our understanding of rapidly evolving organisms or data sets with ancient DNA. Our analyses help us disentangle the definition of these concepts and their practical implications. Our study concentrates on the estimates of the evolutionary rate. Beyond its biological implications, this parameter is essential for estimating evolutionary timescales and our performance metrics for the age of the root-node are comparable to those of the evolutionary rates (see Supplementary material online).

The phylodynamic threshold and measurable evolution are not discrete bounds. Increasing the sampling window generally reduces uncertainty and improves accuracy (bias and coverage), but there is no clear cut-off for when the estimates become accurate and objectively ‘precise’. Notably, the prior on the phylogenetic tree and the evolutionary rate are particularly influential for estimating evolutionary rates and timescales (for a detailed investigation see Tay et al. (2024)).

In fact, data sampled from a population that has attained its phylodynamic threshold does not necessarily mean that the resulting estimates will be correct. In these circumstances the phylodynamic threshold simply means that there is a measurable amount of genetic diversity within the sampling window. However, our measures of coverage, uncertainty and bias for data with sampling windows of 0.5× the expected phylodynamic threshold are often comparable to those obtained with much wider sampling windows, given that the prior on the evolutionary rate is not misleading.

In the case that the prior is substantially biased a sampling window wider than 10× the phylodynamic threshold may be needed to reduce such bias. The extent to which we can draw inferences from data that have attained the phylodynamic threshold or that are measurably evolving depends on the extent to which the data can inform the posterior, which is ultimately a measure of the relative contribution of the prior and the data (via the likelihood function). Our results suggest that using hierarchical priors may be an effective means of minimising the biases due to a poorly chosen prior. We emphasise that a prior that imposes a downward bias appears to be more problematic than one that favours high evolutionary rates. We attribute this to the fact that the upper bound on the evolutionary rate is effectively bracketed by the sampling window. The total genetic divergence divided by the sampling window width constrains the maximum evolutionary rate, whereas in principle there is no bound on the age of a tree and thus the evolutionary rate can effectively reach zero (i.e. if time tends to infinity). We note that the default prior on the evolutionary rate in popular Bayesian phylogenetic packages (e.g. BEAST2 (Bouckaert et al., 2019) and BEASTX (Baele et al., 2025)) is either a uniform distribution between 0 and infinity or a CTMC rate reference prior (Ferreira and Suchard, 2008, Wang and Yang, 2014), both of which tend to favour low evolutionary rate values, and should thus be considered carefully, particularly when the sampling window is narrower than the phylodynamic threshold.

Ideally, the prior and posterior of key parameters, such as the height of the root or the evolutionary rate should overlap, while the posterior should be narrower than the prior (i.e. more informative), meaning that the data and the prior are not in conflict (for recent work on quantifying prior-data conflict see: Nott et al. 2020). In this vein, assessing the adequacy of the model and prior via predictive checks can be illuminating (McElreath, 2018), especially in situations where the joint prior is poorly understood (see Baele and Lemey 2014, Wang and Yang 2014 for discussions about the prior in Bayesian phylogenetics). Recent years have seen the development of a range of methods for assessing phylogenetic model adequacy (Brown and Thomson, 2018, Duchêne et al., 2018, Duchene et al., 2019, McElreath, 2018), for instance one can simulate phylogenetic trees under the posterior estimates to verify whether the height of the root node and the topology could have been generated by the model in question.

Measurably evolving populations are those for which the phylodynamic threshold has been attained and the sampling window is *sufficiently* wide. The criteria for determining the phylodynamic threshold and whether a population is measurably evolving are the same, and are typically assessed via temporal signal. Statistical tests for this purpose quantify the strength of the association between sampling times and genetic distance (Duchêne et al., 2015, Featherstone et al., 2024, Murray et al., 2016, Rambaut et al., 2016, Rieux and Balloux, 2016). That is, the degree to which the sampling times on their own constitute an informative molecular clock calibration. We contend that assessing prior sensitivity is more important than the outcome of tests of temporal signal for obtaining reliable molecular clock estimates. In fact, a poor choice of prior can mislead tests of temporal signal (Tay et al., 2024). If data are drawn from a sampling window that spans the expected phylodynamic threshold, the presence of temporal signal is likely supported by most tests, suggesting accurate estimates. Yet, if the prior used for the estimation is misleading and informative, it might actually obscure the ‘correct’ signal from the data. Thus, positive evidence for temporal signal might not necessarily mean that estimates are accurate. In contrast, if the data are drawn from a narrow sampling window but the prior is reasonable then the estimates may still be reliable, despite a lack of strong temporal signal. It also has to be noted that an increasing sampling effort does not necessarily lead to increasingly correct inferences, because misspecification not only in prior distributions of parameters, but also in the underlying model, can introduce biases (Ferretti et al., 2024, Möller et al., 2018).

An obvious concern about molecular clock calibrations using sequence sampling times is sampling bias. We find that temporal sampling bias, where data are overwhelmingly collected at a particular period of time does not have a substantial impact on estimation accuracy on a simple coalescent model, but that increasing the number of ancient sequences can reduce uncertainty. Another form of sampling bias is when genetic diversity is not uniformly sampled or the underlying population is structured. Previous work has demonstrated that in such cases, the evolutionary rate and the age of the root node tend to be overestimated (Möller et al., 2018), a problem that diminishes when sequence data are increasingly informative or by using a tree prior that explicitly models population structure (e.g. Kühnert et al. 2016, Müller et al. 2017).

Our study has a few limitations that have been partly addressed elsewhere. The number of sequences is fixed in most of our experiments, but it is well known that increasing the number of sequences generally means that data are more informative and thus the estimates are more precise (see Möller et al. (2018) and our Fig.4), and therefore it is likely that the width of the sampling window needed to obtain reliable estimates also depends on the number of sequences. Moreover, our simulation experiments involved a low degree of evolutionary rate variation among lineages. In this respect, it is expected that the width of the sampling window scales with the amount of dispersion in the molecular clock. In addition, our simulations are based on a simple population dynamic model, the constant coalescent. The impact of the sampling scheme and width is likely to be more complex for models with more parameters. We also assume the correct model of evolution and population growth in all simulation-based inferences. With the empirical analysis, we, however, highlight how conclusions drawn from these do not directly extend to real-world data. Rather, the isolated effects found therein describe only one of many elements impacting the inference from real data. Further scrutiny of these factors is warranted, but the main implication is that the necessary sampling time window is a combination of the data set, the organism, and the model at hand.

Overall, our study elucidates some of the fundamental intricacies of molecular clock calibration strategies. We urge researchers to carefully question their model and its underlying prior assumptions, not only via tests of temporal signal, but also through careful choice of the prior, an understanding of the information content in the data, and the implications of model misspecification.

## 4 Materials and methods

### 4.1 Simulations

#### 4.1.1 Data generation

We simulated phylogenetic trees and the evolution of nucleotide sequences to assess the impact of varying the sampling window and temporal sampling bias. We parameterised our simulations to resemble an HBV population evolving over 10,000 years before present, as described by Kocher et al. (2021).

We generated phylogenetic trees under a coalescent process in which the population size has been constant over time using the package ReMaster (Vaughan, 2024), part of the BEAST2 (version 2.7) software. We set the population size to 5,000, which results in trees with an average age of about 10,000 time units (years). The number of samples (i.e. tips) drawn from the trees and their ages were defined in ReMaster, according to the simulation scenario described below.

The simulation of sequence data requires trees with branch lengths in units of subs/site, instead of time. For this purpose, we multiplied the branch lengths of the simulated time-tree with the rate of evolution, a lognormally distributed random variable with parameters 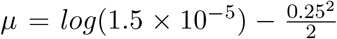 and *σ* = 0.25. This procedure equates to simulating an uncorrelated relaxed molecular clock model with an underlying lognormal distribution (Drummond et al., 2006) with mean, *M* of 1.5 × 10^−5^ subs/site/year and a standard deviation of 0.25 subs/site/year (following that 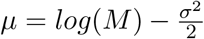). Because we multiply the branch lengths in units of years by a variable in subs/site/year, the resulting trees have branch lengths in units of subs/site, formally known as phylograms, in contrast to time-trees or chronograms where the branch lengths correspond to time. We obtained sequence alignments using the R package phangorn (v2.8.1) (Schliep, 2011), according to a HKY+Γ_4_ substitution model, with parameters *κ* = 2, *α* = 4, and equal base frequencies. The alignments consisted of 3,200 nucleotides to match the average genome size of HBV.

We considered the expected phylodynamic threshold of our data to be about 20 years. For our simulations where we varied the sampling window, we set the ages of 100 tips to be sampled at present (all have an age of 0), or to be drawn from a uniform distribution between 0 and 10 (1/2 of the expected phylodynamic threshold), 0 and 20, 0 and 200, or 0 and 2,000.

To investigate the impact of temporal sampling bias we initially simulated trees with 500 tips with sampling times distributed in 5 time points, with 100 tips per time point. The distribution of sampling times followed an exponential distribution with mean of 4,000, such that sampling times were concentrated towards the present. We simulated these complete trees in ReMaster and we followed the procedure above to simulate a molecular clock model and sequence alignments.

We conducted two sampling schemes for the trees with 500 tips: the ‘time-uniform’ scheme consisted of drawing 20 samples from each time point, whereas the ‘time-biased’ scheme included mostly modern samples (90 from the present, and 5, 3, 1, and 1 from the remaining time points). For each simulation scenario we generated 100 replicates.

#### 4.1.2 Analysis of simulated data

We analysed all data sets in BEAST2 under a model that matched that used to generate the data: the HKY+Γ_4_ substitution model, an uncorrelated relaxed molecular clock model with an underlying lognormal distribution, and a constant size coalescent tree prior. We used the default prior configuration in the program, except for the mean evolutionary rate (*M*) and the population size of the coalescent (*N*, where *N* = *N*_*e*_ × *τ*, and where *N*_*e*_ is the effective population size and *τ* is the generation time). For the population size we assumed *N* ∼ *Exponential*(*mean* = 5, 000), which is centred in the value used to generate the data. The expected height of the root node is roughly 10,000 years (expected time to coalescent=2×*N* for an ultrametric tree (Arbisser et al., 2018, Nordborg, 2019)).

For the mean evolutionary rate we considered a range of priors with different degrees of information content (uncertainty) and for which the mean was either the value used to generate the data, or one order of magnitude higher or lower, as shown in table 6. We also included three degrees of uncertainty in such prior distributions, where the 95% CI width was equal to the mean, or around three or six times as wide. In all cases we set the sampling times for calibration.

**Table 6:**
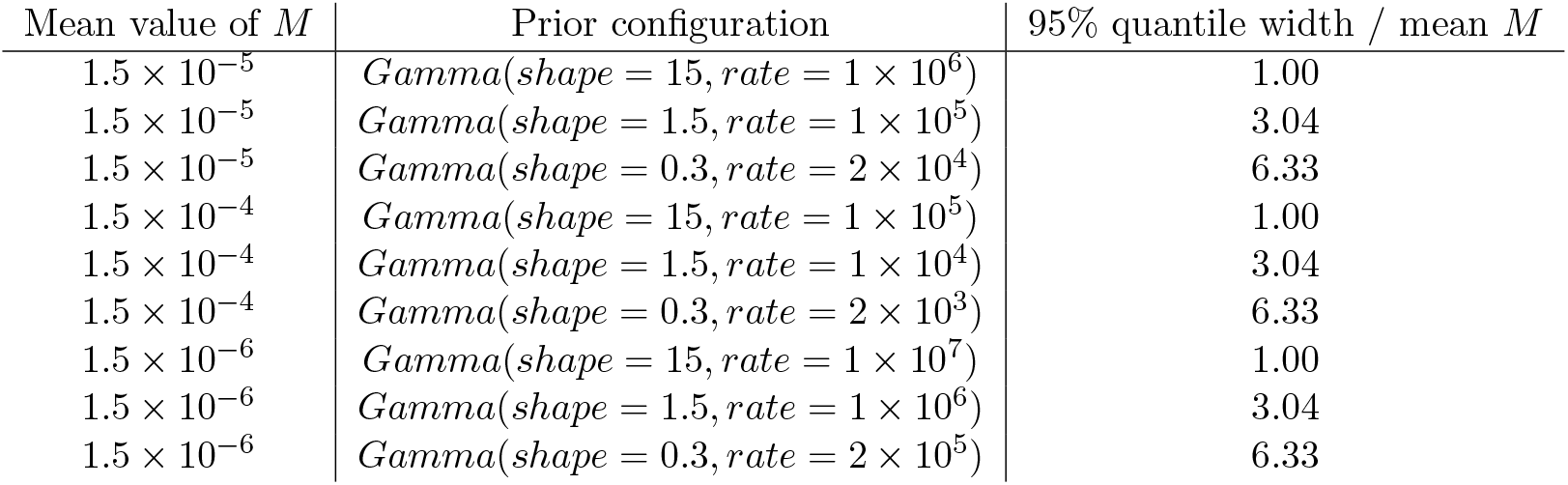
Prior configuration for the mean evolutionary rate, *M* of the lognormal distribution of branch rates. Note that the mean of the *Gamma* distribution here is *shape/rate* and that the true value used to generate the data is 1.5 × 10^−5^ subs/site/year.

For ultrametric trees, sampling times are set to the present, such that all calibration information is provided by the tree prior. Concretely, let the tree length in time (sum of all branch lengths ti in units of time) be *T*_*t*_, the tree height in units of time *T*_*h*_, the branch rates *r*_*i*_ (the vector of branch rates 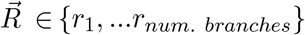, see Douglas et al. 2021), and *T*_*d*_ is the tree length in genetic divergence (sum of branch lengths *d*_*i*_ in subs/site).

In most Bayesian phylogenetic frameworks, branch lengths in genetic distance are the product of rates and times (Douglas et al., 2021, Drummond et al., 2006, Höhna et al., 2016), such that 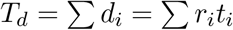. Importantly, *T*_*d*_ is given by the phylogenetic likelihood and only depends on genetic distance and substitution model parameters. The average evolutionary rate (an estimate of parameter *M*) is 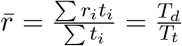. However, *T*_*t*_ is a function of population size *N* through 𝔼 [*T*_*t*_] = 2×*N* ×*log*(*n*), where *n* is the number of sampled lineages (Arbisser et al., 2018, Tavaréet al., 1997). Thus, the prior on *N* affects *T*_*h*_ (through 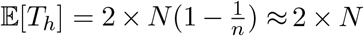, for large *n*), *T*_*t*_, and the average evolutionary rate (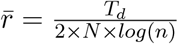, where *T*_*d*_ is independent of time).

We used an additional configuration for the prior *M* using a hierarchical structure as follows:

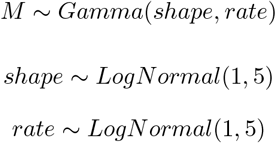

To set up this model, we simply treat the *shape* and *rate* as parameters that are sampled in the model.

We used Markov chain Monte Carlo (MCMC) to sample the posterior distribution for all analyses. We set the chain length to 10^8^ steps, sampling every 5 × 10^4^ steps. We deemed sufficient sampling by verifying that the effective sample size was at least 200, by using the R package CODA (version 0.19) (Plummer et al., 2006). When this criterion was not met we extended the chain length to 5 × 10^8^ steps.

To visualise the prior on *M* in our model with a hierarchical prior we drew MCMC samples from the marginal prior of *M*. That is, the prior integrating over the hyperprior distributions and all other parameters. We obtained such samples by setting the option `sampleFromPrior=“true”` in the input xml files in BEAST2, which conducts the MCMC while ignoring the phylogenetic likelihood.

#### 4.1.3 HBV empirical data

We selected a complete genome data set of HBV published by Kocher et al. (2021). The complete alignment included 232 genomes of length 3,344 nucleotides, with 1,807 variable sites, and 1,498 site patterns. The sampling times ranged from the present to 10,535 years before present. To investigate the impact of varying the sampling window and on temporal sampling bias we subsampled the data as described in our Results section. We analysed each data set using the same model and prior settings as in our simulations.

## 5 Data availability

Computer code, analysis files, and data sets in this study are available at:

https://github.com/sebastianduchene/phylo_threshold_code_data

## 6 Competing interests

None.

## 7 Acknowledgements

We are grateful to the Editor, two anonymous reviewers, and to Sebastian Calvignac-Spencer for valuable comments on earlier versions of this manuscript.

## 8 Funding

This work received funding from the Inception program (Investissement d’Avenir grant ANR-16-CONV-0005 awarded to SD) and the Australian National Health and Medical Research Council (2017284 awarded to SD).

## Notes

https://github.com/sebastianduchene/phylo_threshold_code_data

### Competing Interest Statement

The authors have declared no competing interest.

### Summary of Updates

Additional simulation experiments and novel analyses of empirical data, following a round of revisions

## References

A. Andrades Valtuenã, G. U. Neumann, M. A. Spyrou, L. Musralina, F. Aron, A. Beisenov, A. B. Belinskiy, K. I. Bos, A. Buzhilova, M. Conrad, et al. Stone age yersinia pestis genomes shed light on the early evolution, diversity, and ecology of plague. Proceedings of the National Academy of Sciences, 119(17): e2116722119, 2022.

I. M. Arbisser, E. M. Jewett, and N. A. Rosenberg. On the joint distribution of tree height and tree length under the coalescent. Theoretical population biology, 122:46–56, 2018.

G. Baele and P. Lemey. Bayesian model selection in phylogenetics and genealogy-based population genetics. In M. Chen, K. L, and L. Po, editors, Bayesian phylogenetics, methods, algorithms, and applications, chapter 4, pages 59–93. CPC Press, Boca Raton (Florida), 2014.

G. Baele, X. Ji, G. W. Hassler, J. T. McCrone, Y. Shao, Z. Zhang, A. J. Holbrook, P. Lemey, A. J. Drummond, A. Rambaut, et al. Beast x for bayesian phylogenetic, phylogeographic and phylodynamic inference. Nature Methods, 22(8):1653–1656, 2025.

R. Biek, O. G. Pybus, J. O. Lloyd-Smith, and X. Didelot. Measurably evolving pathogens in the genomic era. Trends in ecology & evolution, 30(6):306–313, 2015.

M. F. Boni, P. Lemey, X. Jiang, T. T.-Y. Lam, B. W. Perry, T. A. Castoe, A. Rambaut, and D. L. Robertson. Evolutionary origins of the sars-cov-2 sarbecovirus lineage responsible for the covid-19 pandemic. Nature microbiology, 5(11):1408–1417, 2020.

R. Bouckaert, T. G. Vaughan, J. Barido-Sottani, S. Duchêne, M. Fourment, A. Gavryushkina, J. Heled, G. Jones, D. Kühnert, N. De Maio, et al. Beast 2.5: An advanced software platform for bayesian evolutionary analysis. PLoS computational biology, 15(4):e1006650, 2019.

L. Bromham, S. Duchêne, X. Hua, A. M. Ritchie, D. A. Duchêne, and S. Y. Ho. Bayesian molecular dating: opening up the black box. Biological Reviews, 93(2):1165–1191, 2018.

J. M. Brown and R. C. Thomson. Evaluating model performance in evolutionary biology. Annual Review of Ecology, Evolution, and Systematics, 49(1):95–114, 2018.

D. A. Buonagurio, S. Nakada, J. D. Parvin, M. Krystal, P. Palese, and W. M. Fitch. Evolution of human influenza a viruses over 50 years: rapid, uniform rate of change in ns gene. Science, 232(4753):980–982, 1986.

A. M. Devault, G. B. Golding, N. Waglechner, J. M. Enk, M. Kuch, J. H. Tien, M. Shi, D. N. Fisman, A. N. Dhody, S. Forrest, et al. Second-pandemic strain of vibrio cholerae from the philadelphia cholera outbreak of 1849. New England Journal of Medicine, 370(4):334–340, 2014.

M. Dos Reis and Z. Yang. The unbearable uncertainty of bayesian divergence time estimation. Journal of Systematics and Evolution, 51(1):30–43, 2013.

J. Douglas, R. Zhang, and R. Bouckaert. Adaptive dating and fast proposals: Revisiting the phylogenetic relaxed clock model. PLoS computational biology, 17(2):e1008322, 2021.

A. Drummond, O. G. Pybus, and A. Rambaut. Inference of viral evolutionary rates from molecular sequences. Adv Parasitol, 54:331–358, 2003a.

A. J. Drummond, O. G. Pybus, A. Rambaut, R. Forsberg, and A. G. Rodrigo. Measurably evolving populations. Trends in ecology & evolution, 18(9):481–488, 2003b.

A. J. Drummond, S. Y. W. Ho, M. J. Phillips, and A. Rambaut. Relaxed phylogenetics and dating with confidence. PLoS Biology, 4(5):e88, 2006.

D. A. Duchêne, S. Duchêne, and S. Y. Ho. Phylomad: efficient assessment of phylogenomic model adequacy. Bioinformatics, 34(13):2300–2301, 2018.

S. Duchêne, R. Lanfear, and S. Y. Ho. The impact of calibration and clock-model choice on molecular estimates of divergence times. Molecular phylogenetics and evolution, 78:277–289, 2014.

S. Duchêne, D. Duchêne, E. C. Holmes, and S. Y. Ho. The performance of the date-randomization test in phylogenetic analyses of time-structured virus data. Molecular Biology and Evolution, 32(7):1895–1906, 2015.

S. Duchene, K. E. Holt, F.-X. Weill, S. Le Hello, J. Hawkey, D. J. Edwards, M. Fourment, and E. C. Holmes. Genome-scale rates of evolutionary change in bacteria. Microbial genomics, 2(11):e000094, 2016.

S. Duchene, D. A. Duchene, J. L. Geoghegan, Z. A. Dyson, J. Hawkey, and K. E. Holt. Inferring demographic parameters in bacterial genomic data using bayesian and hybrid phylogenetic methods. BMC evolutionary biology, 18:1–11, 2018.

S. Duchene, R. Bouckaert, D. A. Duchene, T. Stadler, and A. J. Drummond. Phylodynamic model adequacy using posterior predictive simulations. Systematic biology, 68(2):358–364, 2019.

S. Duchene, L. Featherstone, M. Haritopoulou-Sinanidou, A. Rambaut, P. Lemey, and G. Baele. Temporal signal and the phylodynamic threshold of sars-cov-2. Virus evolution, 6(2):veaa061, 2020a.

S. Duchene, S. Y. Ho, A. G. Carmichael, E. C. Holmes, and H. Poinar. The recovery, interpretation and use of ancient pathogen genomes. Current Biology, 30(19):R1215–R1231, 2020b.

S. Duchene, P. Lemey, T. Stadler, S. Y. Ho, D. A. Duchene, V. Dhanasekaran, and G. Baele. Bayesian evaluation of temporal signal in measurably evolving populations. Molecular Biology and Evolution, 37 (11):3363–3379, 2020c.

K. Eaton, L. Featherstone, S. Duchene, A. G. Carmichael, N. Varlık, G. B. Golding, E. C. Holmes, and H. N. Poinar. Plagued by a cryptic clock: insight and issues from the global phylogeny of yersinia pestis. Communications Biology, 6(1):23, 2023.

V. Eldholm, J. Monteserin, A. Rieux, B. Lopez, B. Sobkowiak, V. Ritacco, and F. Balloux. Four decades of transmission of a multidrug-resistant mycobacterium tuberculosis outbreak strain. Nature communications, 6(1):7119, 2015.

L. A. Featherstone, A. Rambaut, S. Duchene, and W. Wirth. Clockor2: Inferring global and local strict molecular clocks using root-to-tip regression. Systematic biology, 73(3):623–628, 2024.

M. A. Ferreira and M. A. Suchard. Bayesian analysis of elapsed times in continuous-time markov chains. Canadian Journal of Statistics, 36(3):355–368, 2008.

L. Ferretti, T. Golubchik, F. Di Lauro, M. Ghafari, J. Villabona-Arenas, K. E. Atkins, C. Fraser, and M. Hall. Biased estimates of phylogenetic branch lengths resulting from the discretised gamma model of site rate heterogeneity. bioRxiv, pages 2024–08, 2024.

M. Fourment, A. F. Magee, C. Whidden, A. Bilge, F. A. Matsen IV, and V. N. Minin. 19 dubious ways to compute the marginal likelihood of a phylogenetic tree topology. Systematic Biology, 69(2):209–220, 2020.

A. Gavryushkina, T. A. Heath, D. T. Ksepka, T. Stadler, D. Welch, and A. J. Drummond. Bayesian total-evidence dating reveals the recent crown radiation of penguins. Systematic biology, 66(1):57–73, 2017.

M. Ghafari, L. du Plessis, J. Raghwani, S. Bhatt, B. Xu, O. G. Pybus, and A. Katzourakis. Purifying selection determines the short-term time dependency of evolutionary rates in sars-cov-2 and ph1n1 influenza. Molecular Biology and Evolution, 39(2):msac009, 2022.

N. Gharbi, E. Rousseau, and T. Wirth. Clock rates and bayesian evaluation of temporal signal. In Phylogenomics, pages 153–175. Elsevier, 2024.

T. Gojobori, E. N. Moriyama, and M. Kimura. Molecular clock of viral evolution, and the neutral theory. Proceedings of the National Academy of Sciences, 87(24):10015–10018, 1990.

S. Guindon. Rates and rocks: strengths and weaknesses of molecular dating methods. Frontiers in Genetics, 11:526, 2020.

T. A. Heath, J. P. Huelsenbeck, and T. Stadler. The fossilized birth–death process for coherent calibration of divergence-time estimates. Proceedings of the National Academy of Sciences, 111(29):E2957–E2966, 2014.

S. Y. Ho. The changing face of the molecular evolutionary clock. Trends in Ecology & Evolution, 29(9): 496–503, 2014.

S. Y. Ho and S. Duchêne. Molecular-clock methods for estimating evolutionary rates and timescales. Molecular Ecology, 23(24):5947–5965, 2014.

S. Höhna, M. J. Landis, T. A. Heath, B. Boussau, N. Lartillot, B. R. Moore, J. P. Huelsenbeck, and F. Ronquist. Revbayes: Bayesian phylogenetic inference using graphical models and an interactive model-specification language. Systematic biology, 65(4):726–736, 2016.

G. Kahila Bar-Gal, M. J. Kim, A. Klein, D. H. Shin, C. S. Oh, J. W. Kim, T.-H. Kim, S. B. Kim, P. R. Grant, O. Pappo, et al. Tracing hepatitis b virus to the 16th century in a korean mummy. Hepatology, 56 (5):1671–1680, 2012.

A. Kocher, L. Papac, R. Barquera, F. M. Key, M. A. Spyrou, R. Hübler, A. B. Rohrlach, F. Aron, R. Stahl, A. Wissgott, et al. Ten millennia of hepatitis b virus evolution. Science, 374(6564):182–188, 2021.

D. Kühnert, T. Stadler, T. G. Vaughan, and A. J. Drummond. Phylodynamics with migration: a computational framework to quantify population structure from genomic data. Molecular biology and evolution, 33 (8):2102–2116, 2016.

D. Kühnert, M. Coscolla, D. Brites, D. Stucki, J. Metcalfe, L. Fenner, S. Gagneux, and T. Stadler. Tuberculosis outbreak investigation using phylodynamic analysis. Epidemics, 25:47–53, 2018.

S. A. Locarnini, M. Littlejohn, and L. K. Yuen. Origins and evolution of the primate hepatitis b virus. Frontiers in Microbiology, 12:653684, 2021.

R. McElreath. Statistical rethinking: A Bayesian course with examples in R and Stan. Chapman and Hall/CRC, 2018.

F. Menardo, S. Duchêne, D. Brites, and S. Gagneux. The molecular clock of mycobacterium tuberculosis. PLoS pathogens, 15(9):e1008067, 2019.

F. K. Mendes, R. Bouckaert, L. M. Carvalho, and A. J. Drummond. How to validate a bayesian evolutionary model. Systematic Biology, 74(1):158–175, 2025.

M. Merker, J.-P. Rasigade, M. Barbier, H. Cox, S. Feuerriegel, T. A. Kohl, E. Shitikov, K. Klaos, C. Gaudin, R. Antoine, et al. Transcontinental spread and evolution of mycobacterium tuberculosis w148 european/russian clade toward extensively drug resistant tuberculosis. Nature Communications, 13(1):5105, 2022.

S. Möller, L. du Plessis, and T. Stadler. Impact of the tree prior on estimating clock rates during epidemic outbreaks. Proceedings of the National Academy of Sciences, 115(16):4200–4205, 2018.

B. Mühlemann, T. C. Jones, P. d. B. Damgaard, M. E. Allentoft, I. Shevnina, A. Logvin, E. Usmanova, I. P. Panyushkina, B. Boldgiv, T. Bazartseren, et al. Ancient hepatitis b viruses from the bronze age to the medieval period. Nature, 557(7705):418–423, 2018.

N. F. Müller, D. A. Rasmussen, and T. Stadler. The structured coalescent and its approximations. Molecular biology and evolution, 34(11):2970–2981, 2017.

G. G. Murray, F. Wang, E. M. Harrison, G. K. Paterson, A. E. Mather, S. R. Harris, M. A. Holmes, A. Rambaut, and J. J. Welch. The effect of genetic structure on molecular dating and tests for temporal signal. Methods in Ecology and Evolution, 7(1):80–89, 2016.

M. Nordborg. Coalescent theory. Handbook of Statistical Genomics: Two Volume Set, pages 145–30, 2019.

D. J. Nott, X. Wang, M. Evans, and B.-G. Englert. Checking for prior-data conflict using prior-to-posterior divergences. Statistical Science, 35(2):234–253, 2020.

J. Oaks, K. F. Cobb, V. N. Minin, and A. D. Leaché. Marginal likelihoods in phylogenetics: a review of methods and applications. Systematic Biology, 68(5):681–697, 2019.

D. Paraskevis, G. Magiorkinis, E. Magiorkinis, S. Y. Ho, R. Belshaw, J.-P. Allain, and A. Hatzakis. Dating the origin and dispersal of hepatitis b virus infection in humans and primates. Hepatology, 57(3):908–916, 2013.

M. Plummer, N. Best, K. Cowles, K. Vines, et al. Coda: convergence diagnosis and output analysis for mcmc. R news, 6(1):7–11, 2006.

A. Rambaut, T. T. Lam, L. Max Carvalho, and O. G. Pybus. Exploring the temporal structure of heterochronous sequences using tempest (formerly path-o-gen). Virus evolution, 2(1):vew007, 2016.

C. Ramsden, E. C. Holmes, and M. A. Charleston. Hantavirus evolution in relation to its rodent and insectivore hosts: no evidence for codivergence. Molecular biology and evolution, 26(1):143–153, 2009.

A. Rieux and F. Balloux. Inferences from tip-calibrated phylogenies: a review and a practical guide. Molecular ecology, 25(9):1911–1924, 2016.

F. Ronquist, N. Lartillot, and M. J. Phillips. Closing the gap between rocks and clocks using total-evidence dating. Philosophical Transactions of the Royal Society B: Biological Sciences, 371(1699):20150136, 2016.

Z. P. Ross, J. Klunk, G. Fornaciari, V. Giuffra, S. Duchêne, A. T. Duggan, D. Poinar, M. W. Douglas, J.-S. Eden, E. C. Holmes, et al. The paradox of hbv evolution as revealed from a 16th century mummy. PLoS pathogens, 14(1):e1006750, 2018.

R. Sanjuán. From molecular genetics to phylodynamics: evolutionary relevance of mutation rates across viruses. PLoS pathogens, 8(5):e1002685, 2012.

K. P. Schliep. phangorn: phylogenetic analysis in r. Bioinformatics, 27(4):592–593, 2011.

M. A. Spyrou, K. I. Bos, A. Herbig, and J. Krause. Ancient pathogen genomics as an emerging tool for infectious disease research. Nature Reviews Genetics, 20(6):323–340, 2019a.

M. A. Spyrou, M. Keller, R. I. Tukhbatova, C. L. Scheib, E. A. Nelson, A. Andrades Valtuenã, G. U. Neumann, D. Walker, A. Alterauge, N. Carty, et al. Phylogeography of the second plague pandemic revealed through analysis of historical yersinia pestis genomes. Nature communications, 10(1):4470, 2019b.

S. Tavaré, D. J. Balding, R. C. Griffiths, and P. Donnelly. Inferring coalescence times from dna sequence data. Genetics, 145(2):505–518, 1997.

J. H. Tay, A. Kocher, and S. Duchene. Assessing the effect of model specification and prior sensitivity on bayesian tests of temporal signal. PLoS Computational Biology, 20(11):e1012371, 2024.

N. S. Trovão, G. Baele, B. Vrancken, F. Bielejec, M. A. Suchard, D. Fargette, and P. Lemey. Host ecology determines the dispersal patterns of a plant virus. Virus evolution, 1(1):vev016, 2015.

T. G. Vaughan. Remaster: improved phylodynamic simulation for beast 2.7. Bioinformatics, 40(1):btae015, 2024.

B. Vrancken, M. A. Suchard, and P. Lemey. Accurate quantification of within-and between-host hbv evolutionary rates requires explicit transmission chain modelling. Virus Evolution, 3(2):vex028, 2017.

Y. Wang and Z. Yang. Priors in bayesian phylogenetics. Bayesian phylogenetics: methods, algorithms, and applications, pages 5–24, 2014.

R. C. Warnock, Z. Yang, and P. C. Donoghue. Exploring uncertainty in the calibration of the molecular clock. Biology letters, 8(1):156–159, 2012.

G. Zehender, E. Ebranati, E. Gabanelli, C. Sorrentino, A. L. Presti, E. Tanzi, M. Ciccozzi, and M. Galli. Enigmatic origin of hepatitis b virus: an ancient travelling companion or a recent encounter? World Journal of Gastroenterology: WJG, 20(24):7622, 2014.

E. Zuckerkandl and L. Pauling. Evolutionary divergence and convergence in proteins. In Evolving genes and proteins, pages 97–166. Elsevier, 1965.

