## Supplementary material online for "The phylodynamic threshold of measurably evolving populations"

In Fig. 1 we show the posterior estimates of the root height for our simulations of varying sampling window widths. In Fig. 2 we show estimates for varying sampling window width using a uniform prior distribution bounded between zero and infinity.

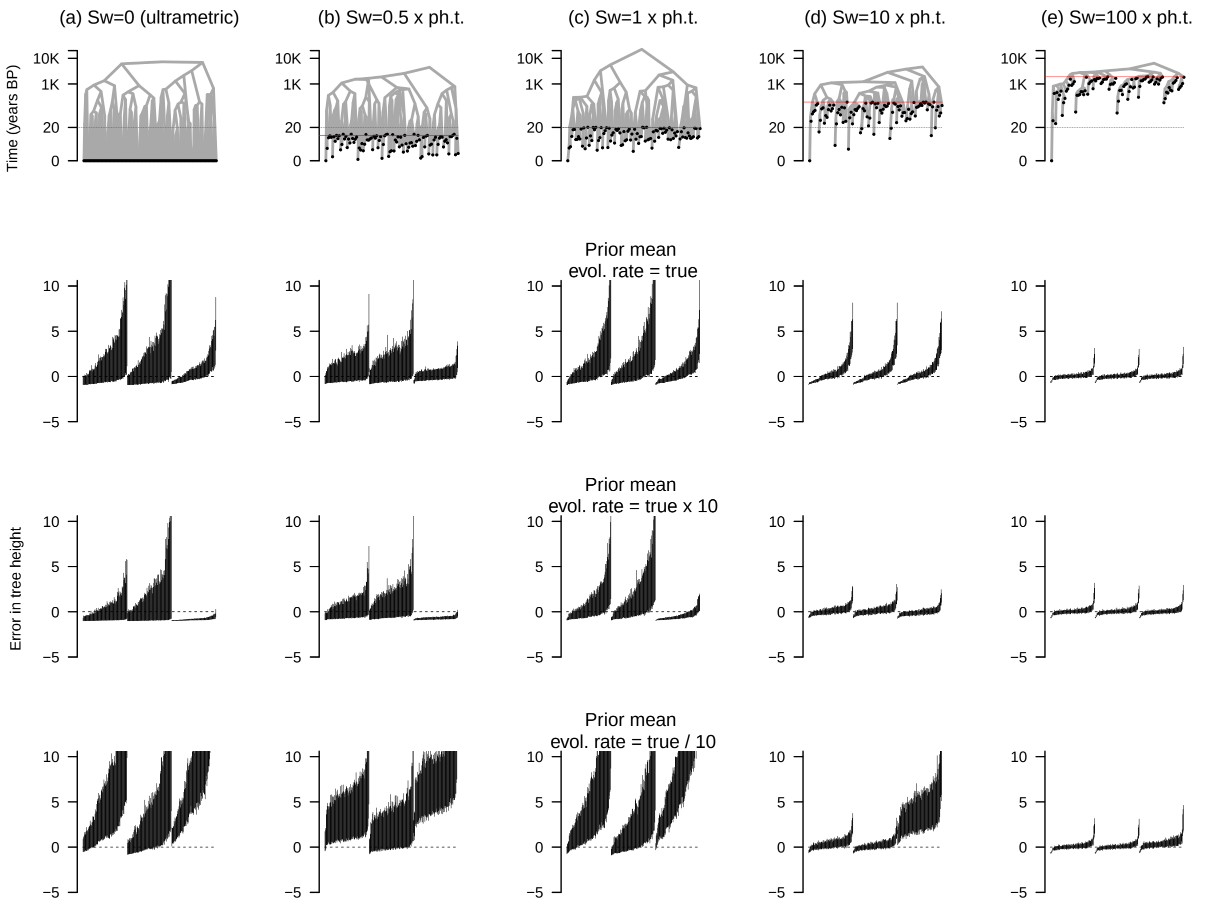


Figure 1: Estimates of the age of the root-node for simulations of varying sampling window widths. Each column corresponds to a simulation setting: (a) is for ultrametric trees where all samples are collected at the same point in time (sampling window, sw=0), (b) is for the situation where the sampling window is 10 years (half the expected phylodynamic threshold; sampling window, sw=0.5× ph.t), (c) is where the sampling window is exactly the expected phylodynamic threshold of 20 years (sw=1× ph.t). Scenarios (d) and (e) denote sampling windows that are 10 and 100 × the expected phylodynamic threshold (sw=10× ph.t and sw=100× ph.t, respectively). The rows denote example phylogenetic trees and prior configurations where the mean evolutionary rate prior, $M$, is set to the correct value (first row), an order of magnitude higher (second row), or an order of magnitude lower (last row). Because each simulated tree has a different root height, we subtract the posterior by the truth and divide by the truth. Thus, a value of 0 is the correct value, whereas one of 1 means a one-fold overestimation. The dashed line is for values of 0. The vertical lines are the 95% posterior credible interval, with 100 replicates for each of the priors on $M$ in table 1.

Table 1: Prior configuration for the mean evolutionary rate, $M$ of the lognormal distribution of branch rates. Note that the mean of the *Gamma* distribution here is $shape/rate$ and that the true value used to generate the data is $1.5 \times 10^{-5}$ subs/site/year.

| Mean value of $M$ | Prior configuration | 95% quantile width / mean $M$ |
|---|---|---|
| $1.5 \times 10^{-5}$ | $Gamma(shape = 1.5, rate = 1 \times 10^5)$ | 3.04 |
| $1.5 \times 10^{-5}$ | $Gamma(shape = 0.3, rate = 2 \times 10^4)$ | 6.33 |
| $1.5 \times 10^{-5}$ | $Gamma(shape = 15, rate = 1 \times 10^6)$ | 1.00 |
| $1.5 \times 10^{-4}$ | $Gamma(shape = 1.5, rate = 1 \times 10^4)$ | 3.04 |
| $1.5 \times 10^{-4}$ | $Gamma(shape = 0.3, rate = 2 \times 10^3)$ | 6.33 |
| $1.5 \times 10^{-4}$ | $Gamma(shape = 15, rate = 1 \times 10^5)$ | 1.00 |
| $1.5 \times 10^{-6}$ | $Gamma(shape = 1.5, rate = 1 \times 10^6)$ | 3.04 |
| $1.5 \times 10^{-6}$ | $Gamma(shape = 0.3, rate = 2 \times 10^5)$ | 6.33 |
| $1.5 \times 10^{-6}$ | $Gamma(shape = 15, rate = 1 \times 10^7)$ | 1.00 |

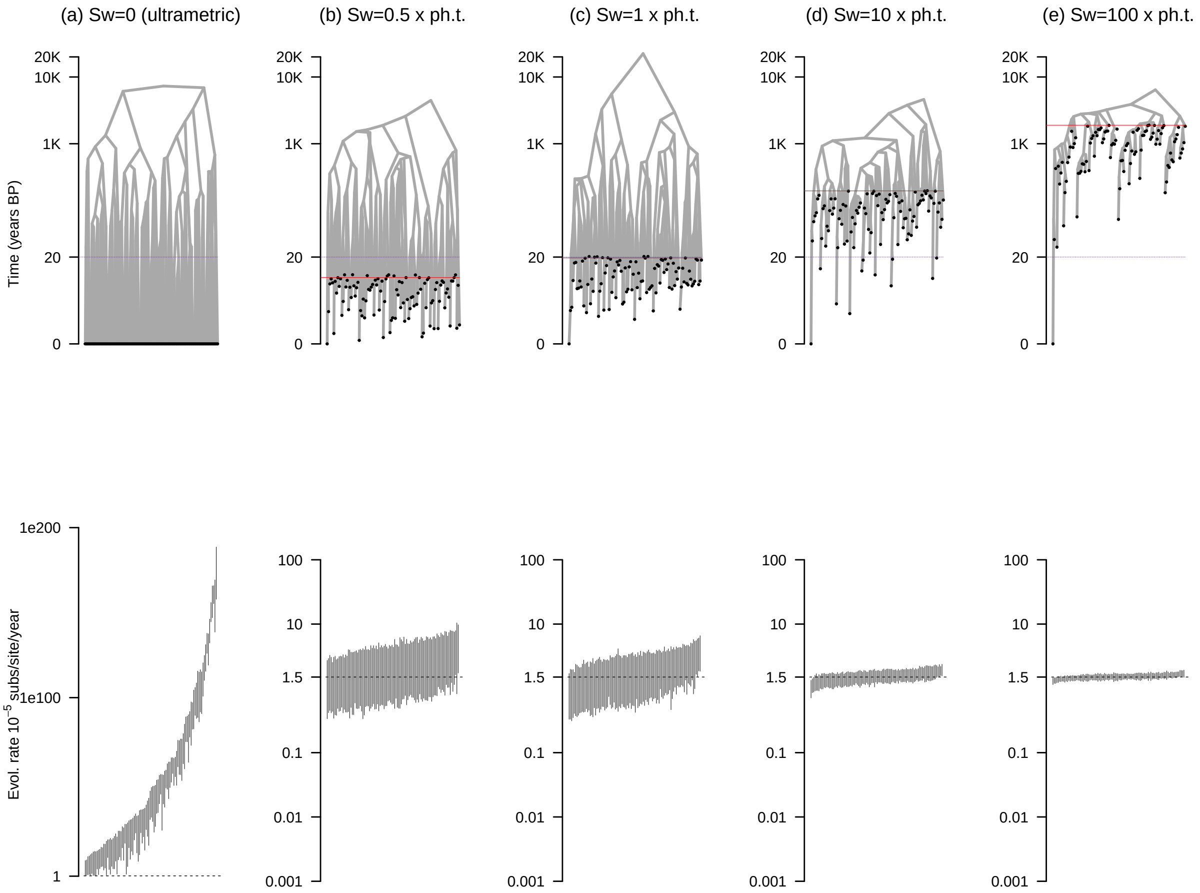


Figure 2: Estimates of evolutionary rates, $M$ for simulations of varying sampling window widths under a uniform prior for $M$. The prior here consists of a uniform distribution bounded between 0 and infinity. The columns match those in Fig. 1. The lines in the second row denote the 95% posterior credible interval across 100 simulation replicates in each case. The dashed lines denote the correct value used to generate the data. Note that the panel in the first column, second row has a different scale for the y-axis.
